# The openretina Project: Collaborative Retina Modelling Across Datasets and Species

**DOI:** 10.1101/2025.03.07.642012

**Authors:** Federico D’Agostino, Thomas Zenkel, Baptiste Lorenzi, Michaela Vystrčilová, Dominic Gonschorek, Samuel Suhai, Samuele Virgili, Alexander S. Ecker, Olivier Marre, Larissa Höfling, Thomas Euler, Matthias Bethge

**Author notes:** These authors contributed equally to this work (Federico D’Agostino, Thomas Zenkel). These authors also contributed equally to this work (Larissa Höfling, Thomas Euler, Matthias Bethge).

## Abstract

The retina provides a unique opportunity to develop a complete and precise model of a computational module in the central nervous system. Deep learning has recently vastly advanced efforts towards this goal, yet decades of data, code, and analysis practices remain fragmented between labs — limiting reproducibility, comparison, and cumulative progress. We argue that an open, collaborative modelling ecosystem is now essential to move the field from isolated studies toward a unified, quantitative account of retinal computation. To this end, we present `openretina`, a modular Python package built on PyTorch that provides a standardised framework for training, evaluating, and interpreting neural network models of the retina. The package implements a shared “Core + Readout” model architecture with a reproducible training pipeline, a common data format based on HDF5, unified evaluation metrics, and *in silico* analysis techniques from the literature. In its initial release, `openretina` integrates five publicly available datasets spanning various species and recording modalities. For each dataset, we provide curated preprocessing, standardised data loaders, and pre-trained model checkpoints that serve as reproducible baselines for benchmarking new approaches. We demonstrate the platform’s utility through example use cases: first, a gradient field analysis linking the instability of optimal stimuli to spatial contrast encoding in ON-OFF retinal ganglion cells; second, systematic benchmarking of architectures within and across datasets, revealing that substantial explainable variance remains uncaptured by current models. By making research tools interoperable across laboratories, `openretina` lays the groundwork for closing this gap collectively.

## Introduction

Modelling the retina has been a long-standing challenge in computational and systems neuroscience. Research efforts towards understanding the complexities of retinal image processing have driven advancements in computational modelling techniques, which in turn have enhanced our understanding of the visual system.

Foundational work established mathematical descriptions of retinal function, such as spatiotemporal filtering models (***Enroth-Cugell and Robson, 1966***), which characterized how different cell types process visual information. Later, statistical models such as linear-nonlinear-Poisson (LNP) models (***Chichilnisky, 2001***) and generalized linear models (GLMs, ***Pillow et al., 2008***) provided a more refined, probabilistic framework for predicting retinal responses. These approaches laid the groundwork for modern system identification techniques but remained inherently limited in their ability to capture the full nonlinear dynamics of underlying retinal ganglion cell (RGC) responses (***Victor, 1988***; ***Deny et al., 2017***; ***Meister, 2025***).

A turning point has been the recent adoption of deep learning tools, particularly convolutional neural networks (CNNs). A lot of research in the retina (e.g. ***McIntosh et al., 2016***; ***Batty et al., 2017***; ***Ding et al., 2021***; ***Qiu et al., 2023***; ***Maheswaranathan et al., 2023***; ***Sridhar et al., 2024***; ***Vystrčilová et al., 2024***) and visual cortices (e.g. ***Klindt et al., 2017***; ***Ecker et al., 2018***; ***Cadena et al., 2019***; ***Burg et al., 2021***; ***Cowley et al., 2023***; ***Du et al., 2024***; ***Wang et al., 2025***; ***Willeke et al., 2026***) has gone into employing artificial neural networks (ANNs) to vastly improve the predictive accuracy and expressivity of system identification models. Once trained, these models can further be used to synthesise a variety of stimuli to characterise neurons and neural populations (***Olah et al., 2017***; ***Walker et al., 2019***; ***Bashivan et al., 2019***; ***Ding et al., 2023***; ***Burg et al., 2024***), and to generate functional hypotheses *in silico* (e.g. ***Höfling et al., 2024***; ***Franke et al., 2022***; ***Goldin et al., 2022***).

Unlike classical approaches, however, the performance of deep learning models is highly sensitive to architectural choices, regularisation, and training procedures, making comparability of community-wide practices and standardised benchmarking more relevant issues. In practice, the tools and data for retinal system identification remain fragmented across laboratories, each with its own codebase and data format, and often differing evaluation protocols. This makes it difficult to reproduce results, compare models, or build on existing work, and raises the barrier to entry for researchers without strong machine learning expertise.

To address this, we present `openretina`, a modular and extensible Python package for easier training, evaluation, and interpretation of ANN-based computational models of the retina.

Our framework builds upon deep learning toolkits for neuroscience, such as `neuralpredictors` (***Willeke et al., 2023c, 2022***; ***Turishcheva et al., 2024b***), on which we base part of our codebase, and `torch-deep-retina` (***McIntosh et al., 2016***; ***Maheswaranathan et al., 2023***), which also provides data-driven models of the retina. Compared to these earlier efforts, with `openretina` we offer a unified, end-to-end solution designed to facilitate model benchmarking and community-driven development, easy dataloading, and model interpretation.

`openretina` directly integrates multiple publicly available retinal datasets (Table 1), making them immediately usable for training neural networks on a wide range of neural responses across organisms (Fig. 1a), recording modalities (Fig. 1b), and stimuli (Fig. 1c). Additionally, the package includes pre-trained model checkpoints for each dataset, allowing users to quickly apply these models without any retraining.

**Table 1.** Overview of datasets currently integrated in `openretina` . Dataset names are with respect to our [huggingface](https://huggingface.co/datasets/open-retina/open-retina) repository. Abbreviations: WN = white noise, MEA = multielectrode array. The reported number of cells is after each dataset’s own quality checks, but prior to possible exclusions based on the Fraction of Explainable Variance (FEV).

| Dataset | Species | Recording Modality | Stimulus Composition | Stimulus Type(s) | Sessions | Cells | Test repeats | Pixel Size [$\mu\text{m}$] |
| --- | --- | --- | --- | --- | --- | --- | --- | --- |
| *Maheswaranathan et al. (2023)* | Tiger salamander | MEA | Greyscale | Jittered natural scenes | 3 | 50 | 5-10 | 50 |
| *Höfling et al. (2024)* | Mouse | 2P $\text{Ca}^{2+}$ imaging | UV / Green | Natural movies | 67 | $\approx 3,000$ | 3 | 12.5 |
| *Karamanlis et al. (2024)* | Mouse & Marmoset | MEA | Greyscale | Jittered natural scenes + WN | 12 & 6 | $\approx 1,800$ & 42 | 18-55 | 7.5 |
| *Sridhar et al. (2024)* | Marmoset | MEA | Greyscale | Natural movies + WN | 4 | 180 | 10 | 7.5 |
| *Goldin et al. (2022)* | Mouse & Axolotl | MEA | Greyscale | Static natural scenes | 1 & 2 | 41 & 50 | 30 & 20 | 42 & 54 |

**Figure 1.**
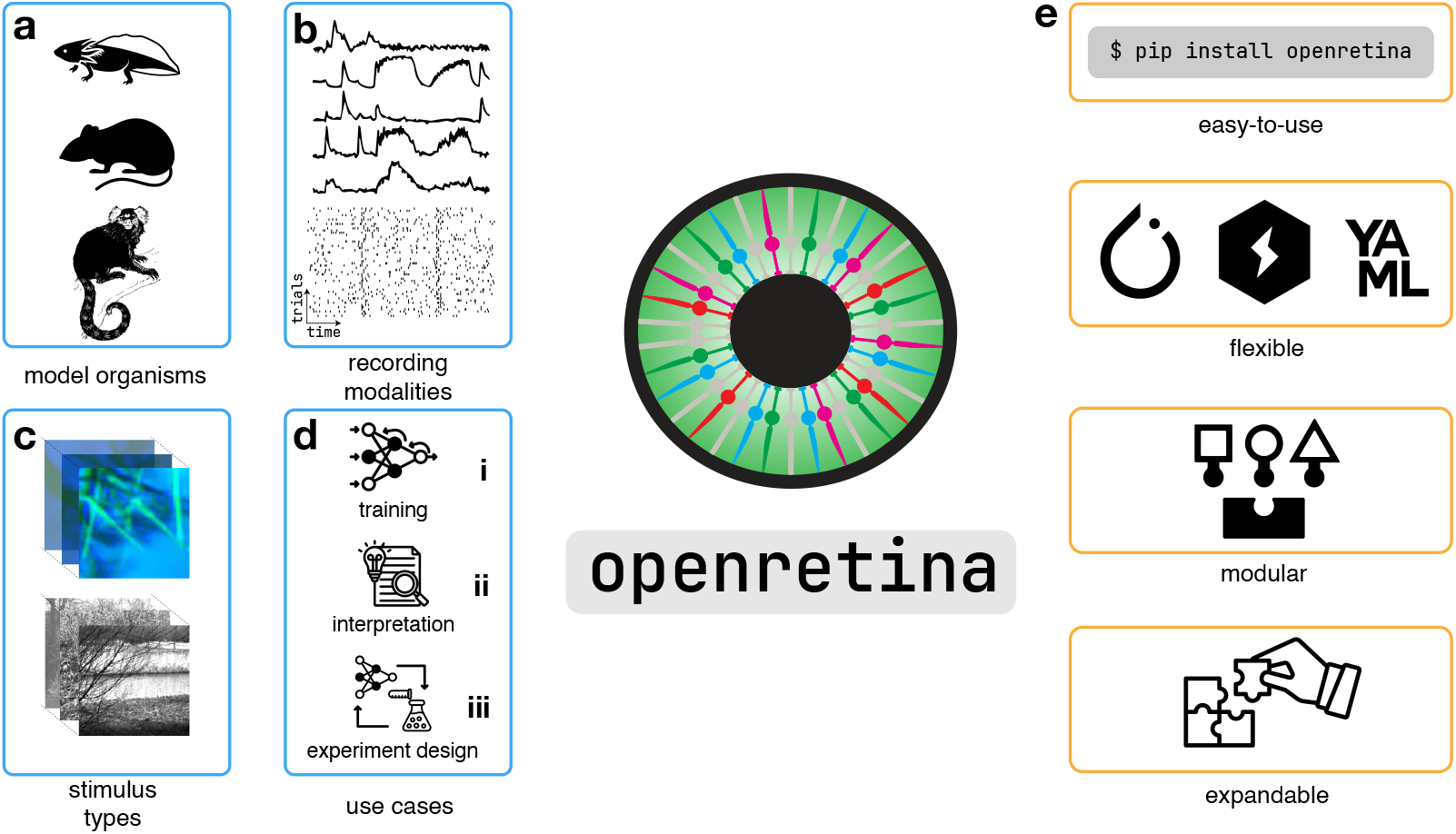
openretina - a Python package for ANN-based computational models of the vertebrate retina. **(a)** `openretina` supports modelling of a variety of experimental data across different model organisms, **(b)** recording modalities (e.g. electrophysiological or imaging recordings), and **(c)** stimuli (e.g. colour and greyscale, images and movies). **(d)** `openretina` allows researchers to use existing models or train their own (**(i)**); to use models for *in silico* testing of hypotheses about retinal computations (**(ii)**), and guiding the design of new experiments (**(iii)**). **(e)** The package is designed to be: easy to use, offering an accessible entry point into retina modelling; flexible, using modern deep learning tools; modular, featuring model architectures that allow combining different components (Core + Readout); and expandable, allowing researchers to contribute new datasets and models, and increasing the value of `openretina` as a resource for the retina research community.

**Figure 2.**
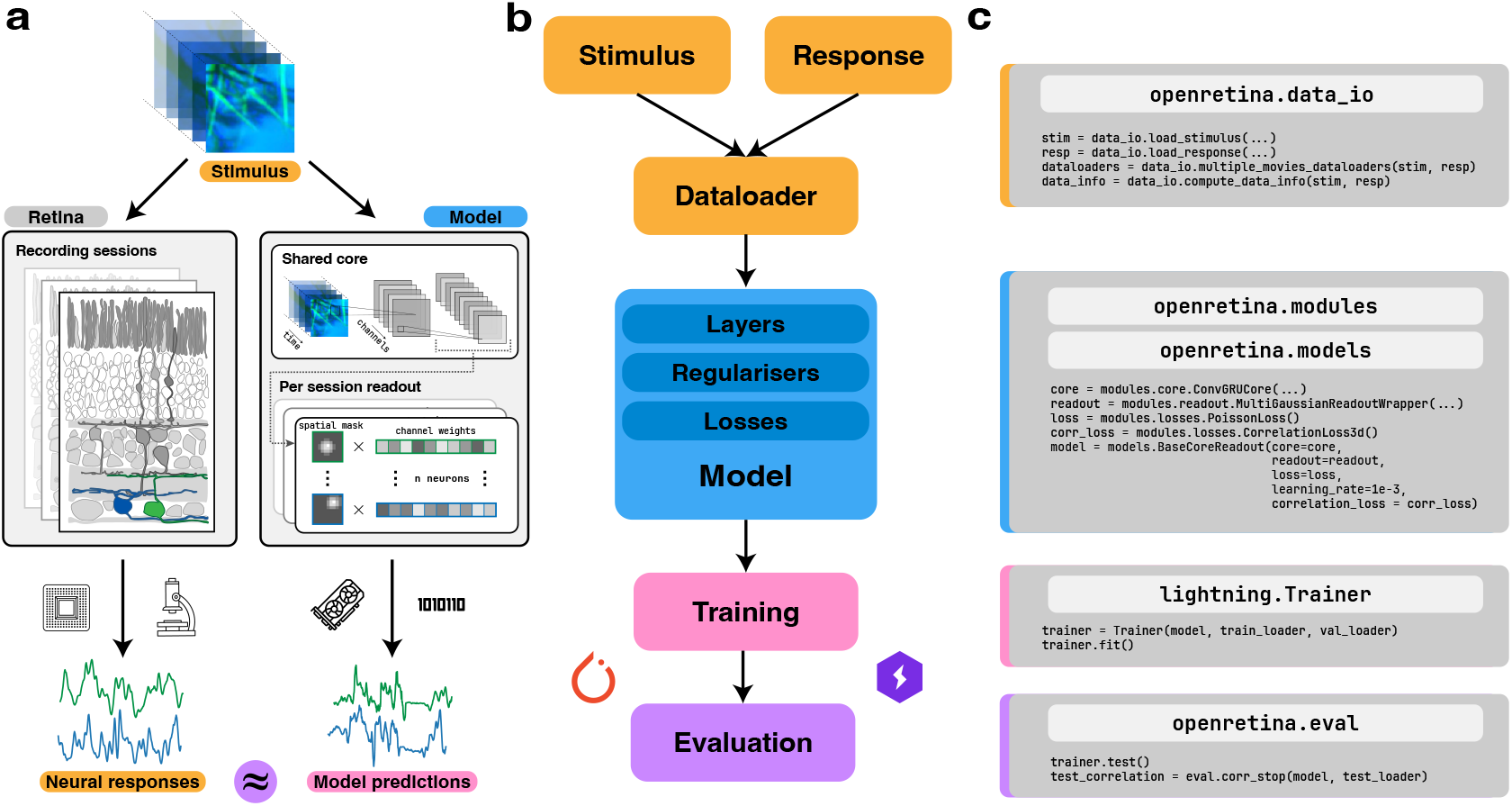
Overview of modelling framework and training pipeline in openretina . (**a**) Typical retina experiments involve recordings of neural responses across multiple experimental sessions. Each session includes paired stimulus and response data, such as videos presented to the retina and the corresponding neural activity recorded via multielectrode arrays (MEAs) or two-photon calcium imaging. Predictive models are then trained using the same stimulus-response pairs to reproduce retinal output. (**b**) Model training workflow. Input data to the model (in orange) for different sessions is processed, split into different folds for training and evaluation, and composed into dataloaders by functions in openretina.data^_^io . The core modelling stage (in blue) passes these inputs into a full model ( `openretina.models` ) composed of layers, regularisers, and loss functions ( `openretina.modules` ), with training managed by PyTorch and Lightning (in pink). After training, evaluation utilities ( `openretina.eval`, in purple) provide metrics, perform comparisons to ground truth, and support testing on novel stimuli. (**c**) Mock training code examples for the different stages. For more detailed explanations and working examples, refer to the package repository and online documentation (open-retina.org/package_docs).

`openretina` also includes visualisation tools for dataset exploration and *in silico* analysis tools to probe and interpret learned representations. Users can visualise model weights, synthesise maximally exciting inputs (MEIs, ***Walker et al., 2019***; ***Bashivan et al., 2019***) that reveal feature selectivity, and extend these optimisation objectives to explore response properties in greater detail (Fig. 1d, ***Goldin et al., 2022***; ***Burg et al., 2024***).

Ultimately, we envision `openretina` as a central hub for neural network modelling of retinal data, offering a growing collection of datasets and tools for model development that bridges across research laboratories. We welcome collaboration and contributions, and are committed to providing an intuitive interface for both machine learning researchers and neuroscientists.

In this paper, we introduce `openretina` as a stand-alone Python package (***D’Agostino et al., 2025***), describing the standardised framework that underpins collaborative modelling and presenting applications that illustrate how it can bridge analysis across laboratories.

Our accompanying website (open-retina.org) provides documentation, tutorials, and dataset cards that complement the material presented here, including step-by-step Jupyter notebooks for different analyses.

## Methods

### Datasets and Models

#### The “Core + Readout” architecture

Models in `openretina` adopt the widely used “Core + Readout” framework (***Klindt et al., 2017***; ***Sinz et al., 2018***; ***Cadena et al., 2019***; ***Ecker et al., 2018***; ***Willeke et al., 2023b***; ***Lurz et al., 2021***; ***Turishcheva et al., 2024a***; ***Ustyuzhaninov et al., 2022***; ***Wang et al., 2025***, among others). This approach separates the model architecture into two distinct components. The *Core*, which is usually shared across neurons, processes the input stimulus extracting a rich feature representation of the visual input. Following this, the neuron-specific *Readout* maps the Core’s output feature space to individual neurons’ firing rate predictions for a given stimulus.

This modularity makes the framework both general and extensible. At one end, classical single-cell Linear-Nonlinear-Poisson (LNP) models can be recast within this setting, with the Core serving as a trivial pass-through and the computational load handled entirely by the Readout. At the other end, any architecture from the broader computer vision community can serve as the Core, provided it maps visual inputs to a spatial feature representation that the Readout can index into, including recurrent architectures such as Gated Recurrent Units (GRU, ***Cho et al., 2014***) and, potentially, pre-trained vision backbones for task-driven modelling (***Cadena et al., 2019***). All models currently implemented in `openretina` are data-driven, meaning both Core and Readout are trained from scratch on stimulus-response pairs.

This decoupling also lays the groundwork for future explorations, such as joint training across heterogeneous datasets or transfer learning across experimental conditions. Practically, this architectural philosophy is implemented in the `openretina.models` and `openretina.modules` subpackages, which enforce the separation of cores and readouts through base classes while offering a library of pre-configured components that users can instantiate directly or combine into custom architectures.

#### A common data format for interoperability

To facilitate collaboration across laboratories with varying experimental setups, the data format used in `openretina` relies only on having access to synchronised stimulus-response pairs. Regardless of the underlying recording modality, data must be structured as time-series inputs (stimuli) and outputs (neural responses). While the native sampling rates of these streams often differ, all `openretina` requires is for them to be temporally aligned or resampled to a common time base prior to modelling, ensuring compatibility with diverse recording modalities.

Following the standard for modern system identification (e.g., ***Maheswaranathan et al., 2023***; ***Willeke et al., 2022***), `openretina` implements image-computable models. Consequently, all stimuli are represented as video tensors rather than parametric descriptions, irrespective of whether the underlying stimulus is white noise, natural video, or synthetic patterns.

To ensure data remains self-describing and portable, we adopt the Hierarchical Data Format (HDF5; ***The HDF Group, 1997-2026***) as the standard storage container. HDF5 allows for the bundling of high-dimensional stimulus and response arrays alongside extensive metadata, which is essential for biological interpretation. This can include experimental covariates such as temperature or adaptation state, preprocessing details like spike-sorting specifics and quality metrics, and functional annotations such as cell-type classifications.

Finally, to promote reproducible benchmarking, we rely on datasets that provide explicit training and test splits encoded in the file structure. Many such datasets also include repeated presentations of the same stimuli in the test set, which enables estimation of trial-to-trial variability and the computation of noise-aware metrics such as the Fraction of Explainable Variance Explained (FEVE; ***Cadena et al., 2019***; ***Pospisil and Bair, 2021***).

To assist users in navigating data resources, Table 1 provides a high-level overview of the datasets currently available in the `openretina` format. Complementing this, the online documentation features “dataset cards”—additional information for each entry. These cards are designed to be kept up to date and evolve alongside the platform, part of a living registry that will expand as new datasets are contributed to the collaborative effort. This granular information is intended to guide researchers in selecting the most appropriate data to answer specific research questions, ensuring that model training is informed by the functional and experimental nuances of the underlying recordings.

On the software side, the openretina.data^_^io module serves as the bridge to the available datasets. It provides utilities for data loading, management of train/test splits, and template classes for including new datasets.

#### Metrics for evaluation and benchmarking

To complete our modelling framework, we provide a unified suite of evaluation metrics and loss functions, as part of `openretina.eval` and `openretina.modules.losses`, respectively. For model optimisation, we support standard objectives such as the Poisson loss, Mean Squared Error (MSE), and Correlation loss. For performance reporting, we also include metrics that account for biological variability. These include, for example, the Fraction of Explainable Variance Explained (FEVE), implemented following ***Cadena et al. (2019*)**, which measures the proportion of stimulus-driven variance captured by the model. As another example, we include a Leave-One-Out (or jack-knife) oracle. This method estimates the theoretical upper bound of model performance on a particular test data set by predicting the response to a specific trial using the average response of all other repeats of the same stimulus, thereby accounting for trial-to-trial variability.

Heterogeneity in experimental protocols—such as varying test set durations, different numbers of stimulus repeats, and diverse preprocessing pipelines—limits unbiased quantitative comparisons across datasets. Despite this, by using a shared codebase for evaluation across datasets, we ensure that reported scores are methodologically consistent, and encourage them to be accompanied by all relevant context (e.g., see Table 2).

**Table 2.** Within-dataset model comparison. Comparison of different model architectures trained on the euler^_^lab/hoefling^_^2024 dataset (***Höfling et al., 2024***) evaluated on the test data. We report Pearson Correlation and Fraction of Explainable Variance Explained (FEVE) across neurons. Correlation is reported for both comparing the model prediction with the average response across test trials (Avg) and to individual trials (Indiv) with ± specifying the standard deviation of the correlation across the individual trials. All models also share the same “time lag”, i.e., the same number of frames consumed by the convolutional kernel along the temporal dimension. All evaluated model assets and hyperparameters are available on [huggingface](https://huggingface.co/datasets/open-retina/open-retina/tree/main/model_checkpoints/2026-03-09).

| Model Architecture | Correlation ↑ | | FEVE ↑ |
| --- | --- | --- | --- |
|  | Avg | Indiv |  |
| LNP | 0.293 | 0.235 (± 0.020) | 0.029 |
| Greyscale Low-Res CNN | 0.550 | 0.438 (± 0.014) | 0.386 |
| Low-Res CNN | 0.549 | 0.438 (± 0.018) | 0.413 |
| Low-Res GRU | 0.585 | 0.466 (± 0.018) | 0.422 |
| High-Res CNN | 0.578 | 0.460 (± 0.018) | 0.450 |
| High-Res GRU | 0.587 | 0.468 (± 0.018) | 0.445 |

#### Unified training and configuration

While the architectural, data, and evaluation standards define the theoretical framework, their practical utility is realised through a unified training pipeline. In `openretina`, we integrate these components into a cohesive workflow that abstracts away the complexity of the training loop, allowing researchers to focus on hypothesis testing.

This unification is achieved technically through a configuration management system (based on ***Yadan, 2019***). By decoupling the model specification from the training logic, users can dynamically swap architectural components or datasets simply by modifying a configuration file or commandline argument.

Under the hood, the training infrastructure leverages PyTorch Lightning (***Falcon and The PyTorch Lightning team, 2019***) to handle essential but repetitive tasks such as logging, checkpointing, and distributed training. The result of this pipeline is a standardised model asset comprising the trained weights and the full configuration state, which serves as an object for all subsequent analysis. It is on this model asset that we can run the diverse *in silico* experiments and applications described in the following sections.

### *In Silico* analyses

Trained predictive networks can be used to investigate the neural computations underlying the recorded responses. The *in silico* techniques present in `openretina` leverage two key advantages of this computational approach, namely differentiability of the trained models and full access to models’ internals, which allow them to be probed in ways that are often difficult or impossible in physiological experiments. As a result, these techniques allow us to probe the strategies learned by the model, generating concrete hypotheses about retinal function that can subsequently be validated *in vivo* and *ex vivo*. The `openretina.insilico` sub-module is designed as an extensible library for these analysis methods.

#### Optimising stimuli

A defining functional feature of a neuron is its optimal stimulus, i. e. the stimulus that drives its activity the most. `openretina` supports estimating this *Most Exciting Input* (MEI, ***Walker et al., 2019***; ***Bashivan et al., 2019***; ***Höfling et al., 2024***; ***Olah et al., 2017***) for RGCs by defining an objective, in this case the average response of a given neuron in a defined time window, and then using the function optimize^_^stimulus to optimise a stimulus towards the given objective. During this optimisation, `openretina` starts with an initialised stimulus (e. g., randomly initialised), and iteratively updates the stimulus in the direction of the gradient to increase the objective.

This optimised stimulus can then be saved as a video, or decomposed into its temporal and spatial components (Fig. 3a; for illustration see the accompanying Jupyter notebook notebooks/mei_example.ipynb).

**Figure 3.**
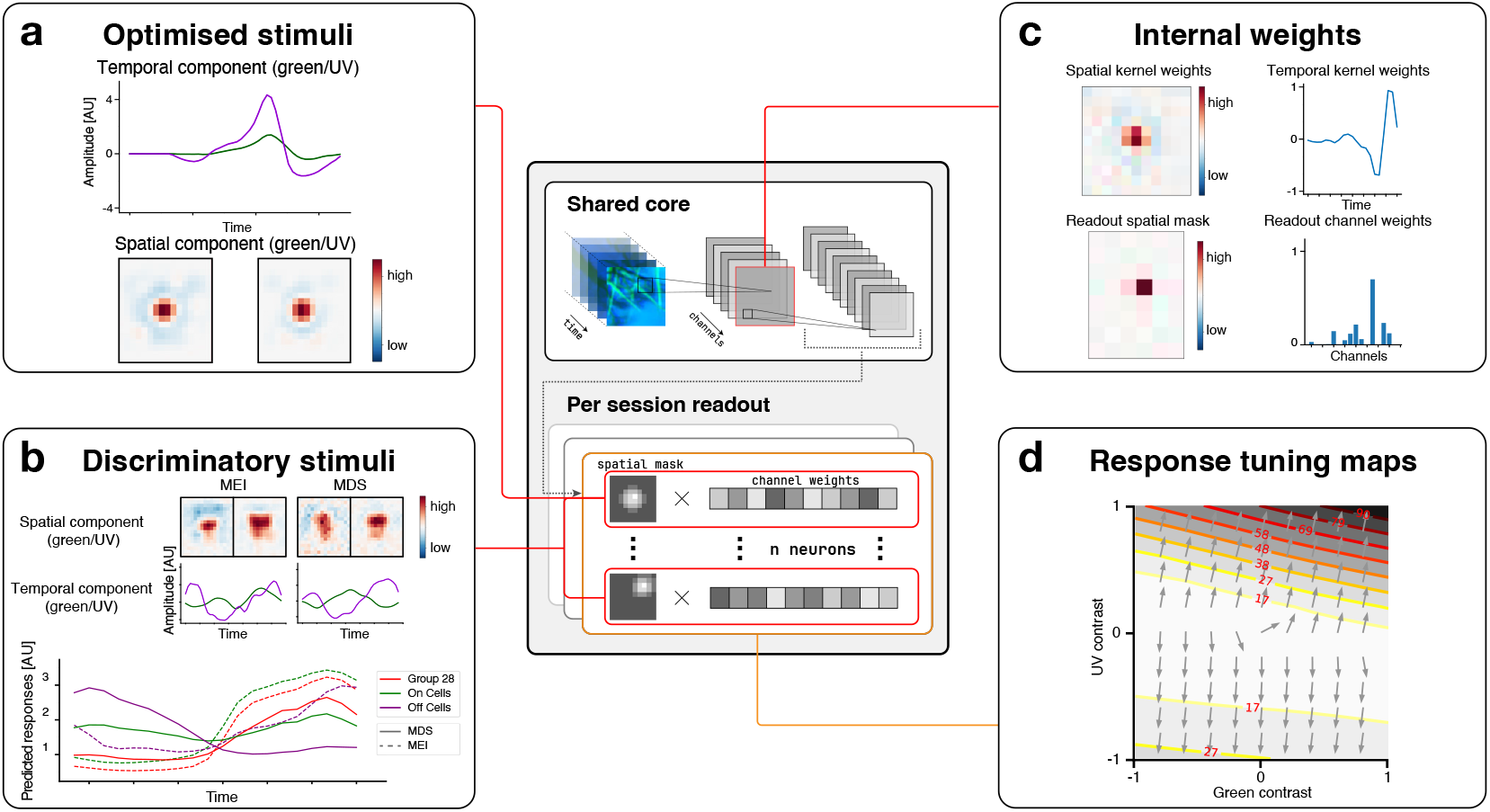
openretina provides tools for visualising model weights and for performing *in-silico* experiments. (**a**) **Optimised stimuli:** The trained model allows us to optimise stimuli towards any objective, e.g. towards activating a particular neuron strongly. The panel shows the spatial and temporal components of an optimised MEI stimulus for a neuron. For the decomposition, we followed ***Höfling et al. (2024***) and ran singular value decomposition on each colour channel of the MEI stimulus individually. The visualisation then shows the first singular vector for the decomposed spatial and temporal dimensions, respectively. (**b**) **Discriminatory stimuli:** Discriminatory objective functions are used to optimise a stimulus towards activating certain neurons strongly while inhibiting other groups of neurons. The upper part of the panel shows both an MEI, optimised to activate all transient Suppressed-by-Contrast RGCs (tSbC) cells, and a most discriminatory stimulus (MDS), which was optimised to drive the activity of tSbC cells, while keeping the activity of ON and OFF cells low. As seen by the average responses in the optimisation window (lower sub-plot), while for the MEI stimulus the average response of ON cells is higher than the response of group 28 cells, the MDS leads to a higher response of group 28 cells compared to other ON cells. (**c**) **Internal weights**: By zooming in and visualising individual model weights, we can better understand how the neural network reproduces the behaviour of the retina. The upper part of the panel shows a visualisation of the spatial and temporal weights of a three-dimensional convolutional layer. The lower part of the panel shows the readout mask, which determines which spatial location in the stimulus influences a modelled neuron, and the readout channel weights that determine the influence of each convolutional channel on the modelled neuron. (**d**) **Response tuning maps:** We can evaluate the model neurons along directions in stimulus space that we are interested in by changing stimulus parameters corresponding to those directions and examining model responses. In this example, we examined the directions along which green and UV temporal contrast selectivities, respectively, change from negative to positive. Lines represent iso-response curves (***Gollisch and Herz, 2012***), arrows are response gradient directions.

To give an example of a more complex optimisation objective in our repository, we show how to generate stimuli that elicit maximally distinct responses for different groups of cells (see visualisation in Fig. 3b). ***Burg et al. (2024***) used this approach to cluster neurons based on their functional responses. Since it is possible to define arbitrarily complex objectives as long as they are expressible as differentiable functions, this provides a powerful interface for probing the neural network (Fig. 3b; notebooks/most_discriminative_stimulus.ipynb).

#### Visualisation and interpretation of model weights

An MEI gives insights about the message that a retinal neuron sends to the brain (***Lettvin et al., 1959***; ***Szatko and Franke, 2022***; ***Reinhard and Münch, 2021***), but it remains agnostic about the retinal circuitry that implements the extraction of this message from visual inputs (***Karamanlis et al., 2022***; ***Kerschensteiner, 2022***; ***Maheswaranathan et al., 2023***). Towards such a mechanistic understanding, we can extend the approach of optimising stimuli to internal “neurons” of the model. However, transferring insights from internal model neurons to biological interneurons requires that the model’s architecture is deliberately designed to reflect the structure of the retinal circuit, for example, in the number of processing stages or the correspondence between convolutional channels and cell types, and that this correspondence is carefully validated. When these conditions are met, interpreting internal representations in biological terms becomes feasible (e. g., ***Maheswaranathan et al., 2023***; ***Schröder et al., 2020***). For the default “Core + Readout” architecture, we support weight visualisation for both the convolutional layer and the readout layer (Fig. 3c).

For convolutional layers, we visualise the weights of each channel of a convolutional layer separately. The spatiotemporally separable convolution layer defined in `STSeparableBatchConv3d` consists of a two-dimensional spatial weight and a one-dimensional temporal weight. The outer product of the spatial weight and the temporal component yields a three-dimensional tensor for the convolution operation.

The Readout layer defined in `SimpleSpatialXFeature3d` can be visualised as a two-dimensional Gaussian mask that defines how different spatial locations in the Core’s feature maps are weighted to predict an RGC response. Similarly, the feature weights that indicate the importance of each channel of the Core for the output of the current neuron are visualised as a one-dimensional bar chart. By combining the visualisations of the weights and the MEIs of the internal neurons, we gain insights into how the neural network calculates its features.

#### Analysing tuning around the optimal stimulus

To move beyond the limited view of a neuron’s response function provided by a point estimates such as the MEI, and to gain a more complete picture of a cell’s tuning such as its selectivity for particular stimuli or its sensitivity to variations along different stimulus dimensions - we can locally expand the tuning curve around its maximum (i.e., the MEI). `openretina` supports this by mapping a model neuron’s response and its gradient around the MEI along user-defined stimulus dimensions.

We illustrate this using the chromatic contrast tuning analysis introduced by ***Höfling et al. (2024***) for an example cell from the same dataset (Fig. 3d). Starting from a neuron’s MEI, we systematically rescale green and UV contrast independently to sample a 2D grid of stimuli around the optimum. At each grid point, we compute both the model’s predicted response and its gradient with respect to the stimulus. The resulting response tuning map and vector field (Fig. 3d) reveal how the cell’s selectivity changes across chromatic contrast space, indicating whether it maintains a stable preference or shifts nonlinearly with stimulus context. A notebook reproducing the analysis is available at notebooks/hoefling_gradient_analysis.ipynb.

## Results

The framework and techniques presented in the previous sections create a shared ecosystem where data, architectures, and analysis methods become interoperable modules. By decoupling these components, `openretina` allows researchers to move beyond isolated studies and leverage community-wide resources to address complex questions. Here, we present two applications that illustrate the practical benefits of this collaborative approach.

### Gradient fields link unstable MEIs with spatial contrast sensitivity

Maximally exciting inputs (MEIs) can be used to characterise the stimulus features that drive a neuron’s response. They represent stimuli that maximise the response as predicted by the model. However, this maximum may not be unique. For example, in the case of ON-OFF cell types (***Höfling et al., 2024***), the MEI optimisation algorithm generates stimuli with opposite polarities for the same cell depending on initialisation (Fig. 4a,b).

**Figure 4.**
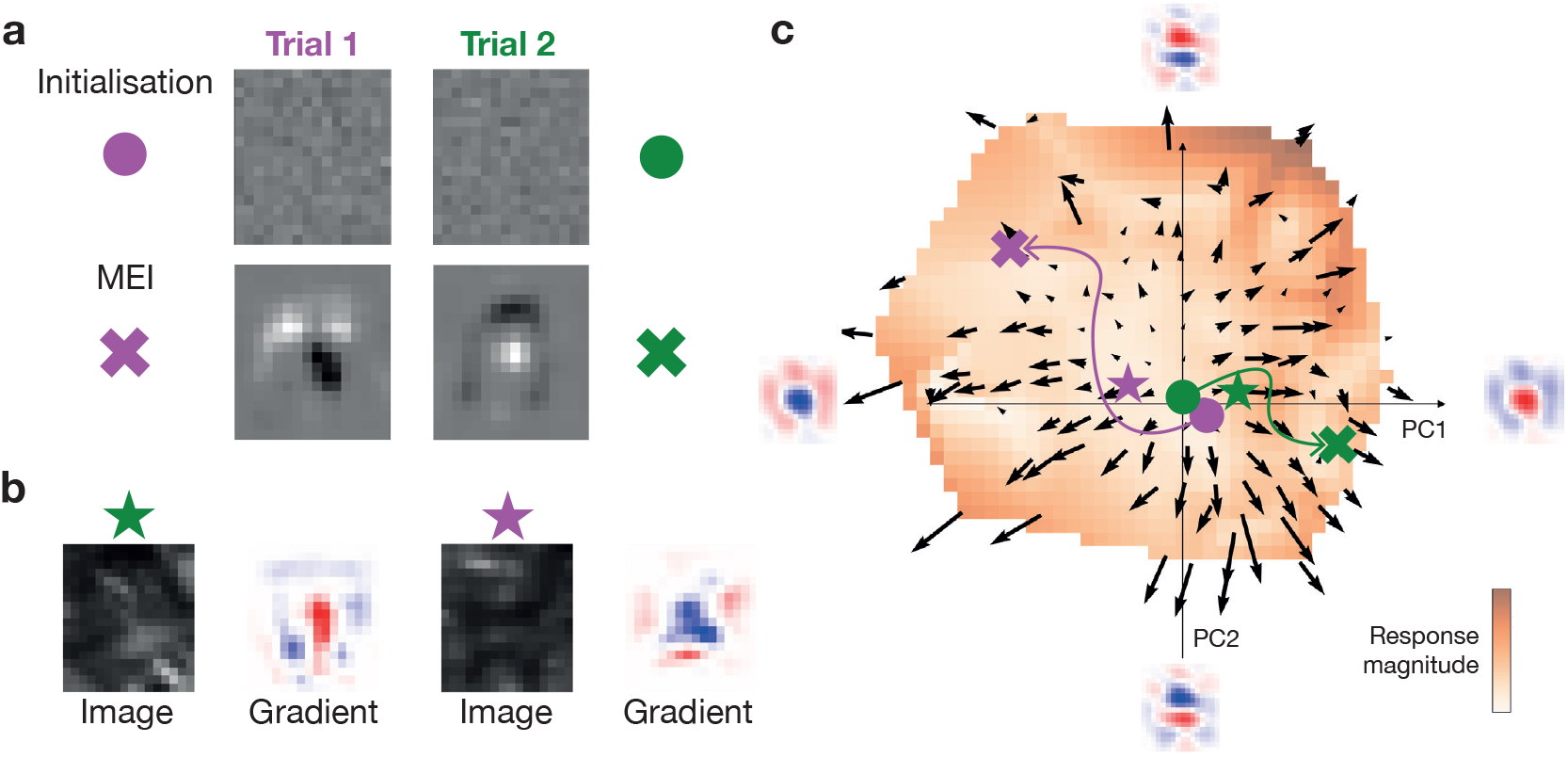
Example application of *in silico* experiments. (**a**) **MEIs:** Green channel participation to the spatial component of the initialisation of MEI optimisation algorithm and associated identified MEI after convergence for an example cell. (**b**) **Local gradients:** Example of image/gradient pairs for two images. This data-driven estimation of the gradient captures the local structure of the cell’s sensitivity in different stimulus context. (**c**) **Gradient field:** Gradient field (green channel) of the same cell. Images in A and B are projected on the vector field and represented using coloured symbols. Arrows sketch potential directions the optimising algorithm could have taken to reach predicted MEIs. Background colour indicates the relative firing rate elicited by stimuli falling in this part of the stimulus space. Insets at the extremes of the axes correspond to the principal components (PCs) (and their negative) of the PCA analysis across all gradients of an example cell.

To understand this phenomenon, we leverage gradients of the stimulus-response function which describe a cell’s sensitivity to changes at each location in stimulus space (Fig. 4a). The gradient acts like a compass for each image, pointing toward the specific visual changes that most effectively boost the cell’s response. We use the model to generate the gradient of the stimulus-response for many different stimuli. We then perform principal component analysis (PCA) on all the gradients, and project both the images and the gradients onto the first two principal components. The arrows in Figure 4c show where to go in the stimulus space to increase the response of the cell; therefore, visualising the gradient vector fields that map a cell’s selectivity across stimulus space (***Goldin et al., 2022, 2023***).

This vector field analysis was applied to a cell classified (***Gonschorek et al., 2025***) as an ON-OFF local edge ‘W3’ RGC (group 10 in ***Baden et al., 2016***) from the ***Höfling et al. (2024)*** dataset. The results offer an explanation in terms of spatial contrast encoding: gradients always point in directions of stimulus space that correspond to increased intensity, regardless of polarity (Fig. 4C).^1^ As illustrated in Figure 4B, MEI optimisation initialisations fall in unstable regions of the gradient field,causing convergence to MEIs with opposite polarities, as predicted for spatial contrast-encoding cells (***Goldin et al., 2022***; ***Virgili and Marre, 2026***).

### Model comparisons

The unified evaluation framework in `openretina` enables systematic model comparisons both within and across datasets. By providing standardised metrics, consistent data splits, and a shared codebase, we allow researchers to assess model performance in a reproducible manner, a prerequisite for identifying which choices genuinely improve predictive accuracy and in which settings, versus which performance gains are artifacts of particular datasets or evaluation protocols. Here, we evaluate models using two complementary metrics. Pearson correlation measures the linear agreement between predicted and recorded responses, capturing temporal structure while remaining invariant to global scaling and offset. The Fraction of Explainable Variance Explained (FEVE) additionally assesses how well the model predicts response amplitudes. For neural responses *y* and predictions ŷ it is defined as

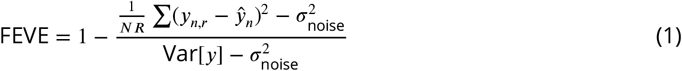

where

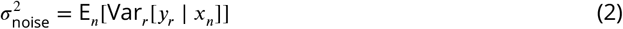

is the trial-to-trial variability averaged across stimuli *x*_*n*_, representing the unexplainable component of the response. FEVE is thus a noise-corrected version of *R*^2^(***Cadena et al., 2024, 2019***; ***Pospisil and Bair, 2021***) and, unlike correlation, is sensitive to systematic scaling errors, conferring more discriminative power (see Suppl. Fig. 5). Following ***Willeke et al. (2023a*)**, we report the correlation both against the trial-averaged response and against individual trials; although the current models in `openretina` produce the same output for each test repeat, the latter allows for future comparisons with models that explicitly capture adaptation effects occurring during multiple trials.

#### Within-dataset comparison

We first benchmark a range of architectures on the test split of the dataset by ***Höfling et al. (2024)***. Across all metrics, neural network-based models, such as convolutional neural networks (CNNs) (***Lecun et al., 1998***) and recurrent neural networks like Gated Recurrent Units (GRU) (***Cho et al., 2014***), outperform linear-nonlinear-Poisson (LNP) models (Table 2). This finding is consistent with results from the Sensorium competition, which benchmarked models on neural recordings from V1 neurons (***Willeke et al., 2023a***).

Switching from greyscale to UV/green video inputs yields a notable improvement in FEVE, while Pearson correlation remains nearly unchanged. Similarly, transitioning from low-resolution inputs (18 × 16 pixels) to high-resolution inputs (72 × 64 pixels) shows a consistent gain in FEVE (see also Suppl. Fig. 5).

#### Cross-dataset comparison

One of the goals of `openretina` is to enable analyses and model comparisons not only within, but also across datasets. However, cross-dataset benchmarking presents substantial methodological challenges. Experimental protocols differ in test set duration, number of stimulus repeats, preprocessing pipelines, and recording quality, all of which influence the measured explainable variance and, consequently, the ceiling against which model performance is evaluated. To partially address this, we adopt a standardised evaluation protocol. Specifically, we restrict analyses to neurons with a fraction of explainable variance above 0.15 (following ***Cadena et al., 2019***), report all metrics using a shared codebase, and document key experimental parameters that affect comparability: most notably, the number of test stimulus repeats, which directly influences the reliability of explainable variance estimates (***Lurz et al., 2022***).

Table 3 reports the results of this evaluation across five datasets. Performance varies considerably: FEVE ranges from 0.449 to 0.710 on models trained on movie inputs, with substantial differences not only between species but also between datasets from the same species and even the same laboratory. Given that these datasets differ along many dimensions simultaneously (species, recording modality, stimulus type, chromatic content, pre-processing, and number of test repeats), attributing performance differences to any single factor is not possible from these comparisons alone. Disentangling these contributions would require systematic, controlled comparisons across a larger collection of datasets than is currently available. Notably, the models trained on ***Goldin et al. (2022)*** achieve markedly higher performance than all other entries; however, these datasets use static natural images with longer presentation periods of each individual image^1^ and simpler, non-temporal model architectures, making direct comparison with the movie-based entries inappropriate. We also note that FEVE estimates are sensitive to the number of test repeats: with as few as three repeats (as in the ***Höfling et al., 2024***, dataset), the explainable variance ceiling is itself a noisy estimate, which can inflate or deflate the resulting FEVE. Entries with fewer test repeats should therefore be interpreted with more caution.

**Table 3.** Cross-dataset model comparison. Evaluation results of pre-trained models provided via `huggingface` . All models use a base, non-recurrent convolutional architecture, and we only include models that were trained on natural scenes or natural movies datasets. To enhance comparability, on all datasets we filter out cells with a FEV lower than 0.15 and report the number of test stimuli repeats, along with different dataset metadata. All models trained on natural movies also share the same “time lag”, i.e., the same number of frames consumed by the convolutional kernel along the temporal dimension. Datasets indicated by “*” use static natural images rather than movies, and the corresponding models operate on single frames without a temporal dimension. All evaluated model assets and hyperparameters are available on [huggingface](https://huggingface.co/datasets/open-retina/open-retina/tree/main/model_checkpoints/2026-03-09).

| Dataset | Specie | Test Repeats | Correlation ↑ | | FEVE ↑ |
| --- | --- | --- | --- | --- | --- |
|  |  |  | Avg | Indiv |  |
| *Maheswaranathan et al. (2023)* | Salamander | 6 | 0.662 | 0.512 ($\pm 0.011$) | 0.528 |
| *Höfling et al. (2024)* | Mouse | 3 | 0.578 | 0.460 ($\pm 0.018$) | 0.449 |
| *Karamanlis et al. (2024)* | Mouse | 32 | 0.724 | 0.442 ($\pm 0.013$) | 0.581 |
| *Karamanlis et al. (2024)* | Marmoset | 32 | 0.814 | 0.448 ($\pm 0.018$) | 0.705 |
| *Sridhar et al. (2024)* | Marmoset | 20 | 0.798 | 0.508 ($\pm 0.041$) | 0.710 |
| *Goldin et al. (2022) \** | Axolotl | 20 | 0.897 | 0.755 ($\pm 0.021$) | 0.813 |
| *Goldin et al. (2022) \** | Mouse | 30 | 0.851 | 0.557 ($\pm 0.049$) | 0.706 |

Nevertheless, two observations are clear. First, current models have yet to reach ceiling performance in prediction: even the best models leave substantial explainable variance unexplained. Second, the fact that performance varies so widely underscores the need for the kind of standardised, multi-lab benchmarking that `openretina` is designed to support: only by evaluating models consistently across a growing collection of datasets can we begin to separate model limitations from data limitations, ultimately advancing our understanding of retinal computation.

## Discussion

With `openretina`, we introduce an accessible Python package designed to train and use ANNs that replicate the input-output behaviour of the retina. By integrating open-source datasets of retinal recordings from mice (***Höfling et al., 2024***; ***Goldin et al., 2022***), salamanders (***Maheswaranathan et al., 2023***; ***Hoshal et al., 2024***; ***Goldin et al., 2022***), and marmosets (***Karamanlis et al., 2024***; ***Sridhar et al., 2024***), `openretina` additionally provides the foundation for a platform for sharing and comparing data and models across research groups, recording modalities and species.

More broadly, we envision `openretina` as a collaborative effort between research groups, and we strongly encourage contributions from researchers and practitioners in the field. Our goal is to create an open, flexible platform that evolves through shared development, and we are open to discussions about design decisions with those interested in shaping the project. Our priority is to build a tool that benefits the field of retinal system identification as a whole.

### Towards collaborative and comparative modelling in visual neuroscience

Platforms for sharing data and comparing models across research groups have gained traction in vision research across species and brain regions. For example, Brain-Score compares normative models of primate ventral stream visual function with respect to their neural and behavioural alignment (***Schrimpf et al., 2018***); the Sensorium Challenge evaluates ANNs of large-scale neuronal activity in the mouse early visual cortex (***Willeke et al., 2023a***; ***Turishcheva et al., 2024b***); and the Algonauts Project (***Gifford et al., 2025***) ranks ANN models of human whole-brain processing of multimodal input. In the field of retinal modelling, ***Vystrčilová et al. (2025)*** provides a systematic comparison of linear-nonlinear models to CNNs on both white noise and natural movie stimuli. These initiatives reflect a shift in computational cognitive and systems neuroscience towards collaborative and comparative modelling and benchmarking. `openretina` presents a first step towards an analogous platform for retinal processing.

### Current limitations and outlook

`openretina` provides a strong foundation for modelling retinal computation, yet several areas remain open for further development: each presenting an opportunity for collaboration and contribution.

Our focus in developing `openretina` so far has been on establishing a flexible and extensible framework, but going forward, we plan to integrate a broader set of models. The package currently focuses on deep neural networks, which have demonstrated excellent predictive performance, but mechanistic models remain essential for understanding the retina’s computational principles (***Schröder et al., 2020***; ***Lappalainen et al., 2024***; ***Deistler et al., 2024***; ***Idrees et al., 2024***). This is where the modular design of `openretina` comes into play. It makes it possible to extend the framework to support also circuit-based or biophysical models. This will then allow to directly compare data-driven and mechanistic approaches, and combine them in hybrid approaches.

To improve comparability between different model classes, we also plan to extend our evaluation methodology and move beyond mean-based evaluations by implementing *Information Gain* (IG) to evaluate full predicted response distributions (***Kümmerer et al., 2015***). To rigorously benchmark this in sparse data regimes (especially in terms of test stimulus repeats), we will adopt the Bayesian oracle estimator (***Lurz et al., 2022***), which generalises the jack-knife approach to provide a robust upper bound on realisable IG.

Another important direction is addressing generalisation across datasets and drawing comparisons across species (***Baden et al., 2020***). Most existing models are trained and evaluated on individual datasets, making it difficult to assess the extent to which learned representations reflect universal principles of retinal computation. Expanding `openretina` to facilitate systematic cross-datasets and cross-species comparisons will help uncover both shared and species-specific processing strategies in the retina.

Finally, a key challenge is the limited availability of openly accessible retinal datasets. For our initiative to reach its full potential as a platform for comparative modelling, more experimental data needs to be shared by the community. We encourage researchers to contribute datasets and collaborate on expanding the available data landscape. In the future, and with the help of contributors, we strive to enrich our dataset records with deeper methodological context, including details on spike sorting practices, stimulus preparation protocols, and other experimental covariates, towards more rigorous modelling practices in the field.

As neuroscience moves toward more open and comparative approaches to modelling, `openretina` aims to be a step in this direction for retinal research. By bringing together datasets, models, and analysis tools in a shared framework, we seek to accelerate scientific discovery and provide a valuable resource for the broader field of computational and systems neuroscience.

## Acknowledgments

We thank Fabian Sinz and the Sinz Lab for developing and open-sourcing `neuralpredictors`, which has been foundational to this work. Key modelling components in `openretina` build directly on their implementations. We are grateful for their commitment to open-science and hope that `openretina` serves as a complementary effort that extends their work and tailors it to the retina community. We thank Jonathan Oesterle for valuable feedback on the manuscript. MB and TE acknowledge support by the Deutsche Forschungsgemeinschaft (DFG, German Research Foundation), CRC 1233 “Robust Vision: Inference Principles and Neural Mechanisms” (TP D2), project number: 276693517. TE acknowledges support by the DFG, EU 42/12-1, project number: 505379160, and iRTG 3130 “limits2vision”, project number: 544833423. MB acknowledges financial support by the Federal Ministry of Education and Research (BMBF), FKZ: 01IS24085B. AE acknowledges support by the European Research Council (ERC) under the European Union’s Horizon Europe research and innovation programme (Grant agreement No. 101041669) and the DFG, CRC 1456, project number: 432680300. LH acknowledges support by the European Research Council through starting grant Eye to Action, 101117156, to Katrin Franke. We thank the International Max Planck Research School for Intelligent Systems (IMPRS-IS) for supporting FD.

## Appendix

**Figure 5.**
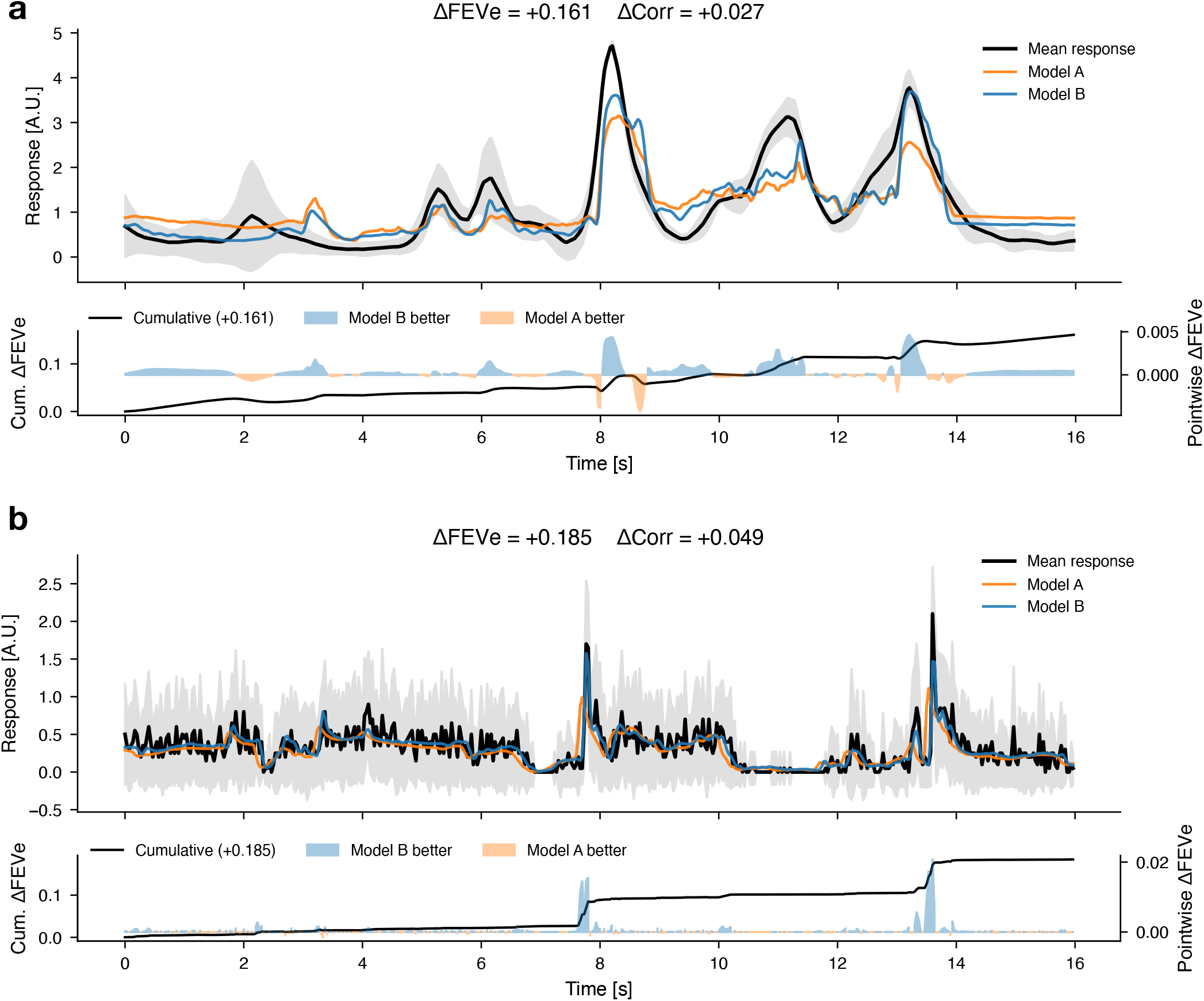
FEVE discriminates between models with the same correlation performance. **(a)** (Top) Predicted responses of Model A (orange) and Model B (blue) for an example neuron from the ***Höfling et al. (2024)*** dataset on a portion of the test set, overlaid on the trial-averaged response (black) and inter-trial variability (shaded, ±1 s.d.). Model B is the high resolution CNN in Table 2, and Model A the low resolution CNN. ΔFEVE and ΔCorr denote the difference in fraction of explainable variance explained and Pearson correlation (Model B − Model A), respectively. (Bottom) Temporal decomposition of the FEVE difference between models. The filled area (right axis) shows the pointwise contribution at each time point, defined as 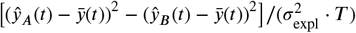, where 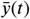 is the trial-averaged response, 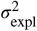 is the explainable variance, and *T* is the number of time points. This normalization ensures the sum over all time points equals Δ FEVE, since 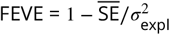 where 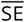 is the temporal mean of the squared errors. Positive values (blue) indicate time points where Model B has lower squared error; negative values (orange) indicate where Model A is better. The black line (left axis) shows the running cumulative sum, which converges to the total ΔFEVE. This decomposition follows from the fact that the noise variance terms cancel when differencing two FEVE values computed on the same neuron (Cadena et al., 2019). **(b)** Same as (a), but on models trained on the ***Sridhar et al. (2024)*** dataset. Model A and B share the same architecture. B is after hyperparameter tuning.

## Footnotes

1 The analysis was performed separately for green and UV channels with equal activations. Natural images were sampled as snapshots from the training dataset.

1 300 ms for individual image presentations in the ***Goldin et al***. (***2022***) dataset compared to a typical frame rate of 30 Hz in movie datasets.

